# Validating a semi-quantitative method to assess the degree of methylene blue staining in sentinel lymph nodes

**DOI:** 10.1101/2020.03.17.994210

**Authors:** Ann Steffi Ram, Michelle L Oblak, Kathy Matuszewska, Ameet Singh, Jim Petrik

## Abstract

**Purpose:** To develop a digital algorithm and validate a semi-quantitative scoring method for surface methylene blue (MB) staining in whole lymph nodes (LN).

**Methods:** Lymph nodes from canine models undergoing sentinel lymph node (SLN) mapping were prospectively assessed *ex vivo* and photographed. Two blinded observers evaluated all images and assigned a semi-quantitative score based on surface staining (0 – no blue stain, 1 – 1-50% stained, 2 – 51-100% stained). A standard reference for degree of blue staining was based on signal-to-background ratios using computer-based imaging software with an output measurement of percentage of staining of the LN. Agreement between observers was assessed using the Kappa coefficient.

**Results:** 124 lymph nodes were included and demonstrated strong agreement (K = 0.8007, p < 0.0001) between results of semi-quantitative scoring and image analysis. Also, strong interobserver and intraobserver agreement was observed for the scoring system (K = 0.8051, p < 0.0001 and K = 0.9493, p < 0.0001, respectively).

**Discussion:** Agreement between the observer-based scoring system and imaging software illustrates a validated method in assessing MB staining, without the need for analysis software. The use of a semi-quantitative scoring system shows promise for a simple, objective assessment of MB staining in surgery and for future study. Lymph nodes can have variable surface colour, which can make assessment of blue staining challenging for novice observers in certain cases. This study describes a digital algorithm for quantitative analysis of blue staining in LN thereby providing a novel and objective reporting mechanism in scientific research involving SLN mapping.

## 1 Introduction

Sentinel lymph nodes (SLN) represent the primary site of solid tumor drainage and are valuable indicators for cancer staging and treatmentrecommendations^1,2^. Detecting SLNs is achieved by using lymphatic tracers, most commonly injected peritumorally, that delineate lymphatic tracts to the sentinel nodes^3^. The gold standard in SLN mapping employs dual tracer techniques involving a combination of radioisotopes, blue dyes and/or fluorescence to increase reliability^4–6^. However, in specific countries and facilities, methylene blue dye is used alone for SLN mapping in light of its cost effectiveness, accessibility and safe outcomes^4,7,8^. Methylene blue is a non-specific blue dye with a good safety profile and has been described for use as an alternative to isosulfan blue and patent blue dye^2,4,9^. Breast cancer studies that employ methylene blue dye alone suggest comparable lymphatic uptake and results to other blue dyes^2,10–14^. One of the challenges associated with the use of methylene blue in SLN mapping, can be the correct identification of methylene blue staining compared to normal surface staining. The discernment of blue stain on a lymph node becomes difficult when clinicians cannot identify whether the discolouration is due to staining or to natural lymph node tissue pigmentations, since lymph nodes are often heterogenous in morphology^15^ and brown tissue can appear as blue. In cases where a dyed lymphatic is not visible, this challenge can post some difficulty to the clinician in confirming that a lymph node is truly sentinel. Ideally, a digital algorithm would be used to depict true staining of tissue in each clinical case, however this is not practical or efficient. Therefore, an objective, accessible method for methylene blue stain quantification on whole tissue is needed to improve reporting.

Mouse models of metastasis represent a commonly used tool to investigate lymph node mapping (Shiro Mori et al., 2013). Oftentimes, mouse strains vary between studies, leading to variability in immune status and resultant LN tissue architecture (Servais, 2011). In addition, it is often difficult to detect lymph nodes in small animal models, and there are practical limitations for gross inspection and imaging. The sub-optimal spatial-temporal differences in mice compared to humans may also pose challenges when investigating the distribution of tracers (Servais, 2011). Alternatively, spontaneously formed cancers in pet dogs often follow similar clinical presentation to humans and mitigate limitations due to size (Gardner, 2015). Imaging techniques between veterinary medicine and human medicine are often similar, easing the translational potential between companion animals and humans (Beer, 2017). In turn, canines are increasingly being used as a pre-clinical model to study human lymphatics (Suami, 2013).

This study uses canine as a pre-clinical model for human cancer to describe the use of a semi-quantitative scoring system and image analysis process for methylene blue stained whole lymph node tissue. The goal is to describe the use of an open-access program for image analysis and validate the semi-quantitative scoring system.

## 2 Methods

### 2.1 Imaging of Lymph Nodes Stained with Methylene Blue

Lymph nodes were obtained consecutively from canine patients undergoing full regional lymph node extirpation as part of another SLN biopsy study at the Ontario Veterinary College Health Sciences Centre from 2017-2019. In all patients, peritumoral or intratumoral methylene blue was injected at a concentration of 0.5mg/mL and routine lymph node extirpation performed. Lymph nodes were imaged in either an unstandardized or standardized fashion, depending on which part of the accrual period they were removed. For the unstandardized group, imaging of lymph nodes was performed with an iPhone 6 or newer model, equipped with a 12 mega-pixel camera (Apple, California, USA) in an unstandardized fashion with no lighting, background or camera distance controls. For the standardized group, lymph nodes were placed on a uniform white background in a photo lightbox (Amazon, Canada) equipped with LED lights to provide optimal imaging conditions for gross specimens^16,17^. Using an iPhone X equipped with a 12-megapixel wide angle camera and secondary telephoto lens (Apple, California, USA) images were taken at a distance of 20cm from the specimen through a 1cm hole at the top of the box to improve focus of the smartphone lens.

### 2.2 Semi-quantitative Scoring of Methylene Blue Stain

All lymph nodes were assessed as positive or negative for methylene blue staining by the PI in situ and immediately following removal and assigned a score based on the coloration of the surface of the lymph node. After data collection was completed, all images for both the unstandardized and standardized groups were randomized for evaluation. The randomized images were evaluated by 2 investigators (M.O., A.R.) for blue stain visualized on the surface of the lymph node. Evaluation consisted of a scoring system based on the amount of methylene blue. The scores were determined as follows: 0 = no blue stain, 1+ = 1-50% of the surface of the lymph node is stained, 2+ = 51-100% of the lymph node is stained.

### 2.3 Verification of Image Analysis Method

The image analysis process was verified by testing true negative and true positive lymph node control images. The negative controls are images of clinically normal lymph nodes from routine biopsies, while the true positive lymph nodes are obtained from cases that show distinct and large amounts of blue stain on the surface of the lymph node, which will be further cropped to areas of stain only; in the form of a region-of-interest (ROI)^18^. Due to the use of a pre-existing, built-in plugin for deconvolution of stains and pigments in an image, the plugin contains verified vectors that correspond to specific stains^19^. The main validation required is if the plugin can detect methylene blue staining on whole tissue specimens and the threshold that disseminates dark blue staining from false-positive signals. Based on the output of the plugin, negative control images were tested to adjust the threshold of “blue” staining^20^. The threshold depicts that dark pixels corresponding to “blue” range from 0 (being the darkest) to 125. Pixels above this threshold were observed to be false signals of blue due to image reflection, image quality or dark tissue. Using a threshold of 0-125 after running the deconvolution plug-in on negative control images depicted 0% of stain detected for all controls. Similarly, positive control pictures that primarily focused on heavily stained regions shows >95% signal detection.

#### Quantification of Methylene Blue Stain Using Image Analysis

Images were analyzed in 6 randomized groups of 25; consisting of 124 images analyzed in total. Images were processed and analyzed in FIJI (National Institutes of Health, Bethesda, Maryland, USA), a distribution of an open-source image processing program (ImageJ)^19^. Colour and background corrections were performed on the true colour (RGB) image using the “Subtract Background” feature and auto adjustment function of the “Brightness/Contrast” tool. The scale was set to “no scale” and, by default, the area was measured in pixels. A ROI was drawn around the whole lymph node using the “Colour Threshold” function, producing an automatic threshold over the image that was measured as the area of the entire surface of the lymph node. If auto threshold did not accurately differentiate the area of the lymph node from the image background, the thresholding brightness and saturation levels were adjusted. Subsequent use of the Colour Deconvolution plugin on the RGB image separated pigments into channels. This plugin is usually applied for the purpose of separating multiple histological stains in a tissue sample^21^. The plugin produces a choice of vectors which are associated with specific dye mixtures. For the purpose of this study, the Gimesa vector was chosen. Gimesa is a dye that contains a mixture of methylene blue, eosin, and an optional third component, such as Azure B^21^. Even though the Gimesa setting is for a combination of three stains, the methylene blue vector is verified and readily isolated^21^. The output of the other channels will not show meaningful signals for stains, such as eosin. Once the plugin runs the command, it produces 3 channels of the image in 8-bit grayscale format. For the purpose of this study, the “Colour_1” window was used since it corresponds to the methylene blue channel. The 8-bit image underwent thresholding (Threshold function) using a threshold of 0-125 and a binary image of the thresholded area was generated.

This is a strict threshold based on controls to only detect dark pixels. This threshold was measured and generated the area of methylene blue stain on the surface of the lymph node. The amount of methylene blue stain on the surface of the node is calculated by:

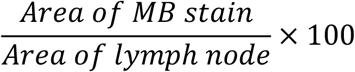

### 2.4 Statistical Analysis

All analyses were completed using SAS 9.3 by a biostatistician (G.M.) who was blinded to the clinical procedures and assessment protocols. Descriptive analysis was performed with summaries of frequencies and agreement percentages in contingency tables. Weighted kappa (κ) statistics were used to assess agreement of assessment modalities, inter- and intra-rater agreement, and agreement between scoring settings. Weighted kappa values with 95% confidence intervals (CI) were calculated to determine strength of agreement. Coefficients in the range of 0.21–0.40 were interpreted as fair agreement, 0.41–0.60 as moderate agreement, 0.61–0.80 as substantial agreement, and 0.81–1 as almost perfect agreement^22^. Statistical significance was set at a two-sided p-value less than 0.05. The sample size is justified using the weighted kappa and degree of agreement.

## Results

A total of 120 lymph nodes were collected from 29 clinical cases of patients undergoing lymph node extirpation with SLN mapping. Eight control lymph node images from 3 clinical cases were also included for analysis. Four images were excluded due to poor image quality or small size of the lymph node resulting in a sample size of 124. Lymph node images were sorted and analysed as described in the methods (Figure 1a). A total of 89 images were included in the unstandardized group and 31 images in the standardized group. The majority of lymph nodes (66.1%) were considered negative for blue staining based on visual assessment, with 29% and 4.8% falling into the 1+ and 2+ categories, respectively. Image analysis was successfully performed and documented in all lymph node images using ImageJ. There was significant concordance between the scoring system and image analysis (κ = 0.8007 [0.70–0.90], P < 0.0001) (Table 1). Interobserver and intraobserver agreement of scoring was strong ((κ = 0.8051 [71–89], P < 0.0001 and κ = 0.9493 [0.89–1], P < 0.0001, respectively). Rater 1 and rater 2 scoring compared to the gold standard displayed similar and substantial agreement (κ = 0.7915, P < 0.0001 and κ = 0.8098, P < 0.0001, respectively). Evaluation of settings based on *ex vivo* scoring and scoring based on images displayed almost perfect agreement (κ = 0.9215 [0.86–0.98], P < 0.0001) (Figure 2).

**Figure 1.**
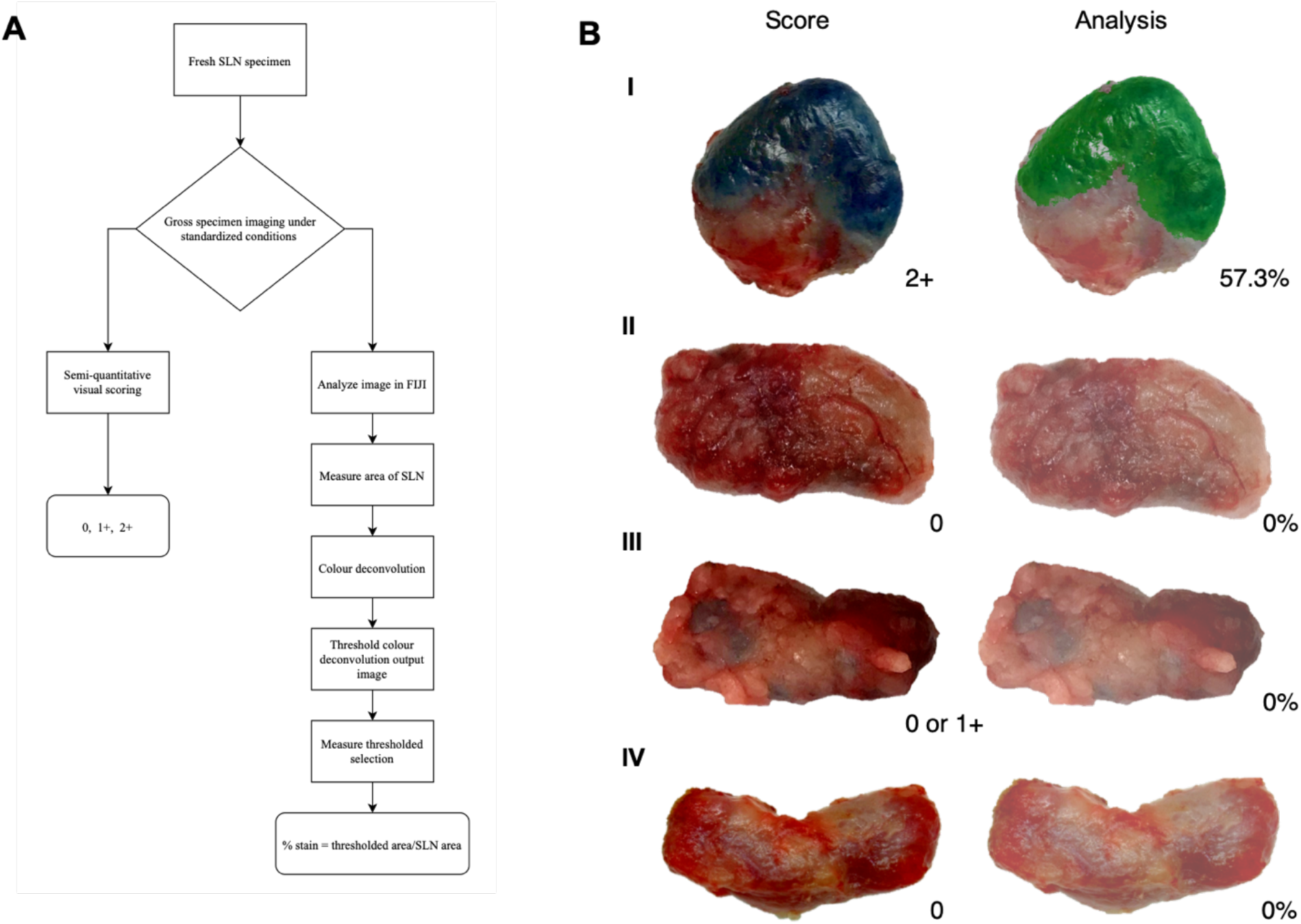
Visual assessment and image analysis outcomes. **a** Workflow of assessment for methylene blue stained lymph nodes. **b** Score and analysis of SLNs. Depicting a positive node (I), a negative node (II), an ambiguous node (III), and a negative control node (IV); all with agreement between score and analysis.

**Table 1.**
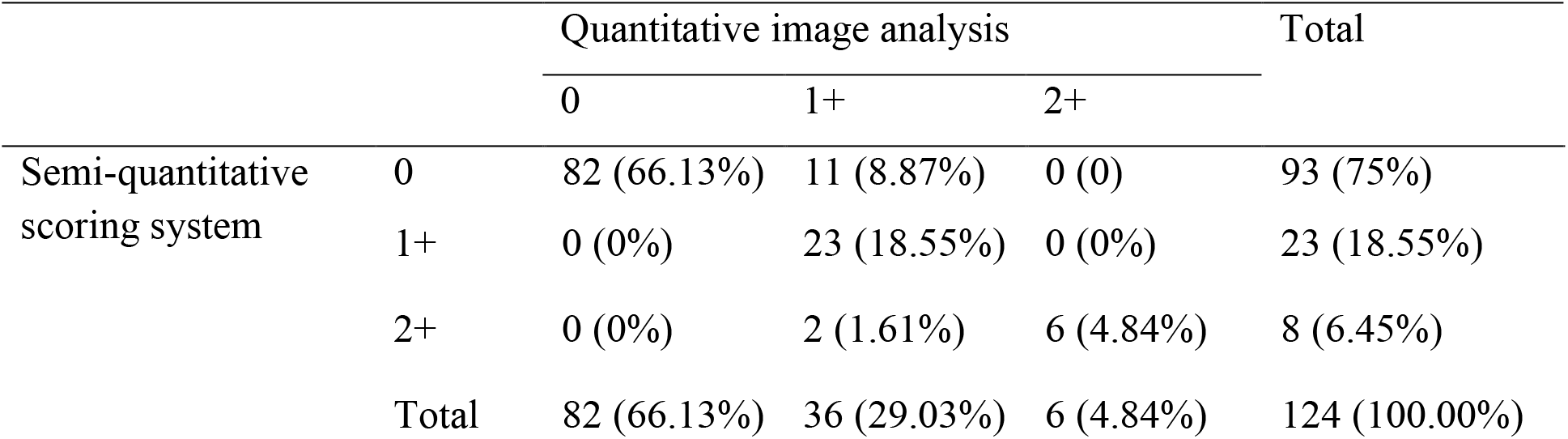
Frequency of agreement between visual scoring and analysis

**Figure 2.**
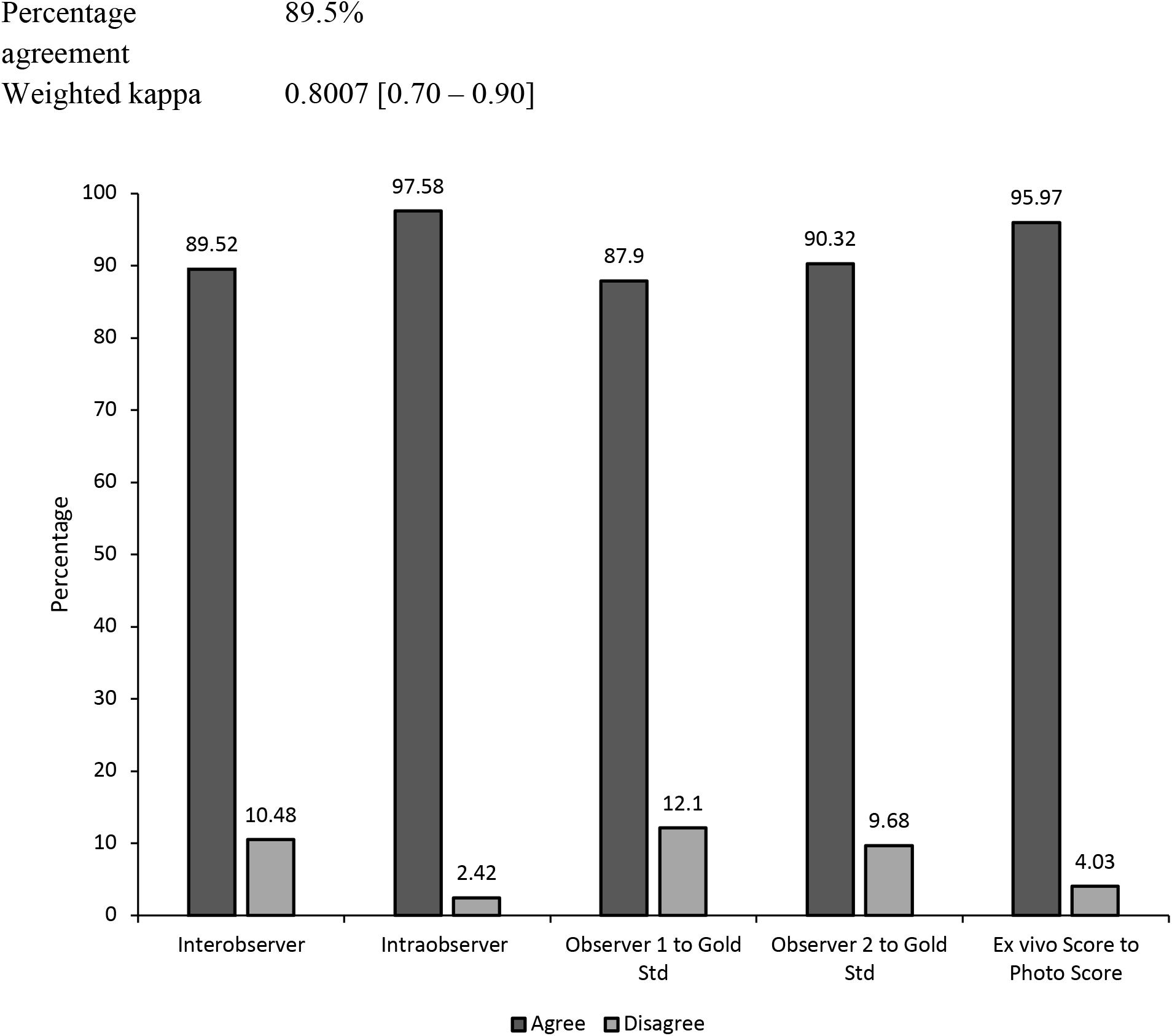
Histogram showing the frequencies of agreement and disagreement of scores in interobserver test, interobserver test, comparing observers to the gold standard, and evaluating observer settings. All frequencies are statistically significant (P < 0.0001).

## Discussion

Sentinel lymph node mapping relies on accurate visual identification of lymph nodes thought to be “sentinel” based on tracer uptake and stains. Literature and research methodologies lack reporting and objective classification of lymph node staining obtained from SLN biopsies. Reports indicate identification rates (IR) as a point of success in the SLN mapping process^4^, although appropriate criteria for successful identification of a SLN is often missing. Clinical trials involving SLN mapping outline inclusion criteria for SLNs reported based on nodes that are only stained blue^2,8,23–28^, blue and non-blue nodes with dye uptake in afferent lymphatic channels^11,12,14,29–35^, or do not comment on IR inclusion criteria^36–38^. The inconsistent or lack of standardized, objective reporting across SLN mapping trials and cases that utilize methylene blue skews accuracy and reduces comparability of results between studies. Improving the reporting process of identified SLNs positive for methylene blue staining can influence the results of studies and enhance discernment of clinicians that assess these nodes.

In this report, we propose a simple, semi-quantitative scoring system to clinically assess methylene blue staining on lymph nodes extirpated during SLN mapping and validated the system using an image analysis process developed for quantification of stain. The assessment of staining is based on the amount of stain present on the surface of lymph node tissue. Our data shows a strong agreement between the scoring system and image analysis that was statistically significant. Immunohistochemistry (IHC) is the gold standard for evaluating stains, however this method is not clinically feasible for postoperative evaluation and is not developed to detect intraoperative methylene blue staining^39^. In contrast, image analysis provides an objective, quick examination of tissue and can be used to improve histology process^40–44^. The image analysis method demonstrated in this study was verified using images of negative control lymph nodes that did not pick up signals for stain and visibly true positive nodes that quantified all the methylene blue staining (Figure 1b). During image analysis, the colour deconvolution (CD) method was employed due to the heterogeneity of lymph node tissue^15,45^. Colour deconvolution allows the separation of RGB colours from images into stain channels made with specific vectors^46,47^. These stain channels are in grayscale and correspond to the intensity of a particular stain found in the image^46^. This analysis plugin determines the density of stain in areas where multiples stains are co-localized^41^. Other studies, such as one by Onder *et al.* (2014) have evaluated the robustness of the CD process and found that CD displayed significantly higher sensitivity in classification of stained samples without compromising specificity when compared to hue-saturation-intensity (HSI) separation method^47^. Due to the superiority of CD in being able to detect dark areas that correspond to brown or blue which HSI could not differentiate, this plugin was employed in our image analysis. The three-grade scoring system developed in this study provides a simple semi-quantitative and accurate assessment of lymph nodes in a clinical setting and is consistent with existing scoring processes that are based on the overall stain intensity (i.e. percentage of cells stained)^40,42,48^. The validation of our scoring system is based on IHC scoring methods that use image analysis to objectively validate the visual assessment^49–51^.

Our scoring system has substantial interobserver agreement depicting there is little variance between scores given by rater 1 and rater 2. The lack of variability between two scorers illustrates that this visual assessment can be done by different observers and still yield the same score given to a sample. Variability in scores between observers is seen when lymph node tissue colouration is ambiguous or if image quality is poor. The intraobserver agreement of scores in our study are near perfect and scores did not even vary when the blinded rater scored the randomized images in batches at a different time. Scores of different observers compared to the gold standard result were in strong agreement, further showing that there is little variability between the score determined by individuals and that of the gold standard. Also, in-person scores of *ex vivo* SLNs were in agreement with scores based on images. This result displays the congruency between samples and images of the samples, where colour and quality are not misconstrued allowing for scoring to take place at any time based on available pictures.

A strength of this study is the high degree of agreement that allows for the sample size to be sufficient with 100% power. The image analysis program is practical and can easily be utilized by clinicians, researchers, and other hospital or lab staff since FIJI is an opensource, free software tailored for biosciences^19^. The accessibility of the program can allow widespread use, allowing for limited variability in analysis programs between studies. Researchers may be less inclined to adopt this method of analysis if an expensive specialized software was required. Also, this visual assessment and image analysis program provides a short learning curve in appropriately scoring and analyzing the lymph nodes. A limitation of this study was the distribution of samples represented in each scoring category. In the concurrent study, patients were undergoing total lymph node basin extirpation to assess the accuracy of SLN mapping in that patient population. As a result, there are a large number of lymph node samples imaged that were negative for methylene blue stain (66.1%). Since the goal of this study was validation, we felt it was important to include this population since it was possible that our visual assessment of a lymph node negative for staining could be different than what was found with image analysis. As a result, our study has an unequal sample size within score categories of 1+ and 2+ (33.8%). However, this does not affect the agreement coefficients due to the analysis and scoring system having low false negative and false positive rates (Table 1). Another potential limitation is the use of CD. Color devolution can have pitfalls in its ability to detect dark areas, where brown pigment can be falsely recognized as dark blue pixels^18,46^, usually when stains like diaminobenzidine (DAB) are used. This was seen in our results where SLNs scored as 0 are detected as a category of 1+ by image analysis (8.87%) since dark tissue is recognized as traces of blue. During our analysis this was not a prominent issue due to the vector used and the strict thresholds. If the program does falsely recognize dark tissue as blue stain, the detection percentage is low and clinically negligible to the human eye where it ranges from 1-2% of stain detected. A final perceived limitation of the study may be the lack of automation to further the objectivity of the analysis process. An automated process for determining ROIs would be efficient and robust, however lymph nodes vary greatly in size and shape which make it difficult to tailor specific macros for ROI creation.

In conclusion, we developed a pragmatic visual and analytic assessment system to evaluate the degree of blue staining in extirpated lymph nodes when SLN mapping is performed using methylene blue dye. The scoring system and quantitative image analysis program have strong agreement which shows the validity of the visual assessment. This assessment workflow allows for standardized reporting of clinical research to improve comparability and consistency of results in SLN mapping of various cancers utilizing methylene blue. The validated visual scoring system provides an accessible and objective measure in a clinical setting when image analysis is not available. It is not yet known if methylene blue staining patterns and uptake has significance for patterns of metastasis and outcomes, but a scoring and digital quantification system is required to investigate such research and is an objective of future directions.

## Acknowledgements

We would like to thank Gabrielle Monteith (University of Guelph) for assisting with statistics used in this report.

## Disclosure of Potential Conflicts of Interest

The authors have no conflicts of interest to disclose.

## Notes

### Competing Interest Statement

The authors have declared no competing interest.

### Summary of Updates

Explanation of the use of canine model

